# Plasma membrane localization of paralogous leucine permeases Bap2 and Bap3 is regulated by Bul1

**DOI:** 10.1101/2020.07.01.181636

**Authors:** S Maheswaran, Paike Jayadeva Bhat

## Abstract

Timeliness in expression and degradation of the nutrient permeases is crucial for any organism. In *Saccharomyces cerevisiae*, post translational regulation of nutrient permeases such as trafficking and turnover are poorly understood. We found that loss of a leucine permease *BAP2*, but not other permeases lead to severe growth retardation when the carbon source is glucose or galactose but not glycerol and lactate. Leucine prototrophy suppressed the retardation, showing *BAP2* and *LEU2* are synthetically lethal. We discovered that loss of *BUL1*, an arrestin involved in trafficking of diverse permeases suppressed this lethality. The suppression required another leucine permease, *BAP3*. Our results suggest that *BUL1* downregulate permeases *BAP2* and *BAP3* present in plasma membrane through Rsp5 dependent endocytosis. We speculate that by regulating leucine import *BUL1* regulates the activity of TORC1.

## Introduction

Leucine is a unique amino acid. It is the highest occurring amino acid in polypeptides and also functions as a signal (Crozier et al., 2005; Stipanuk, 2007; Suryawan et al., 2011; Ananieva et al., 2016). Target of Rapamycin (TOR) the mastermind of cell growth and proliferation in eukaryotes is activated by leucine (Bonfils et al., 2012; Han et al., 2012; Gonzalez and Hall, 2017). Leucine consumption has implications in several diseases. Leucine deprivation results in stalling of breast cancer growth and induces apoptosis (Xiao et al., 2016). Leucine and its breakdown products reduce the glucose uptake, in adult cardiomyocytes even after insulin treatment by affecting translocation of GLUT4, a glucose transporter, establishing a possible link between diabetes and leucine consumption (Renguet et al., 2017).

Despite its ability to make all 20 proteinogenic amino acids, *S.cerevisiae* has amino acid permeases to import amino acids from the external milieu (Ljungdahl and Daigan-Fornier, 2012). These permeases are synthesized in rough endoplasmic reticulum, forwarded to golgi bodies from where they are either dispatched to plasma membrane or vacuole for degradation (Lauwers et al., 2009). When not needed, as in case of absence of substrates, to limit the import in case of excess substrates or while experiencing stress like deviation from optimal temperature, pH, heavy metal ions, improper folding permeases are removed from the plasma membrane through endocytosis, sent to vacuole for degradation (Abe and Iida, 2003; Kim et al., 2007; Lin et al., 2008; Nikko et al., 2008; Nikko and Pelham, 2009; Hatakeyama et al., 2010; O’Donnell et al., 2010; Becuwe et al., 2012a; Zhao et al., 2013; Suzuki et al., 2013; Ghaddar et al., 2014; Crapeau et al., 2014; Talaia et al., 2017; Hovsepian et al., 2017; Hovsepian et al., 2018; Savocco et al., 2019; Tanahashi et al., 2021).

Prior to endocytosis, permeases are ubiquitylated by Rsp5 Ubiquitin ligase with the help of adaptor proteins known as <u>A</u>rrestin <u>R</u>elated <u>T</u>rafficking adaptors (ARTs) also called simply as arrestins (Lin et al., 2008; Belgareh-Touze et al., 2008; Leon and Tsapis., 2009). Although many ubiquitin ligases are known, only the HECT domain Rsp5 ligase, is so far reported to participate in permease ubiquitylation. There are about 10-12 cytoplasmic arrestins and 5-6 membrane bound arrestins in *S.cerevisiae* (Nikko and Pelham, 2009). Each arrestin is responsible for ubiquitylation of a set of permeases and conversely a permease shall have one or more arrestins to facilitate ubiquitylation (O’Donnell and Schmidt, 2019). After ubiquitylation, the permease is endocytosed to form an early endosome, which matures to a late endosome before finally fusing with the vacuole (Rotin et al., 2000; Hettema et al., 2004). This turnover and insertion of newly synthesized permeases in the plasma membrane is necessary for the cell to cope and survive in a constantly changing environment.

Here, we show that the growth of *S.cerevisiae* is critically dependent on *BAP2*, a high affinity leucine permease, either in a synthetic medium containing several amino acids or in a minimal medium with scanty leucine. This observation comes as a surprise considering the ability of several other permeases to transport leucine. We observed that loss of *BUL1*, an arrestin, overcame the dependency on *BAP2*, by stabilizing *BAP3* in the plasma membrane. We show that the turnover of leucine permeases Bap2 and Bap3 is accomplished by Bul1. Finally, we propose that by regulating the abundance of these permeases, Bul1 modulates the activity of TORC1, which in turn regulates Bul1, thus setting up a feedback loop. Hence, this study for the first time provides a possible mechanism as how leucine regulates its own uptake.

## Results

### BAP2 and LEU2 are synthetically lethal in carbon source dependent manner

Commonly used wild type strains (WT) of BY4741, W303 and ∑1278b background are leucine auxotrophs (*leu2*Δ) and suffer a growth defect in a modified SCGlu medium (<u>S</u>ynthetic <u>C</u>omplete <u>Glu</u>cose medium) (Cohen and Engelberg, 2007). The growth defect is due to impaired leucine uptake, suppressed by multicopy plasmids either bearing *LEU2*, *BAP2* or *TAT1* (Cohen and Engelberg, 2007). We believe that the impairment is due to a fierce competition between leucine and other surplus competing amino acids for their import through a common transporter(s). In such a case, reduced abundance of the competing amino acids would favor leucine import and restore growth. Likewise, WT BY4741 grows well in the routine SCGlu medium, which has only 12 amino acids in lower concentrations (Fig.1A), opposed to all 20 amino acids in high concentrations in the modified SCGlu medium.

**Figure 1.**
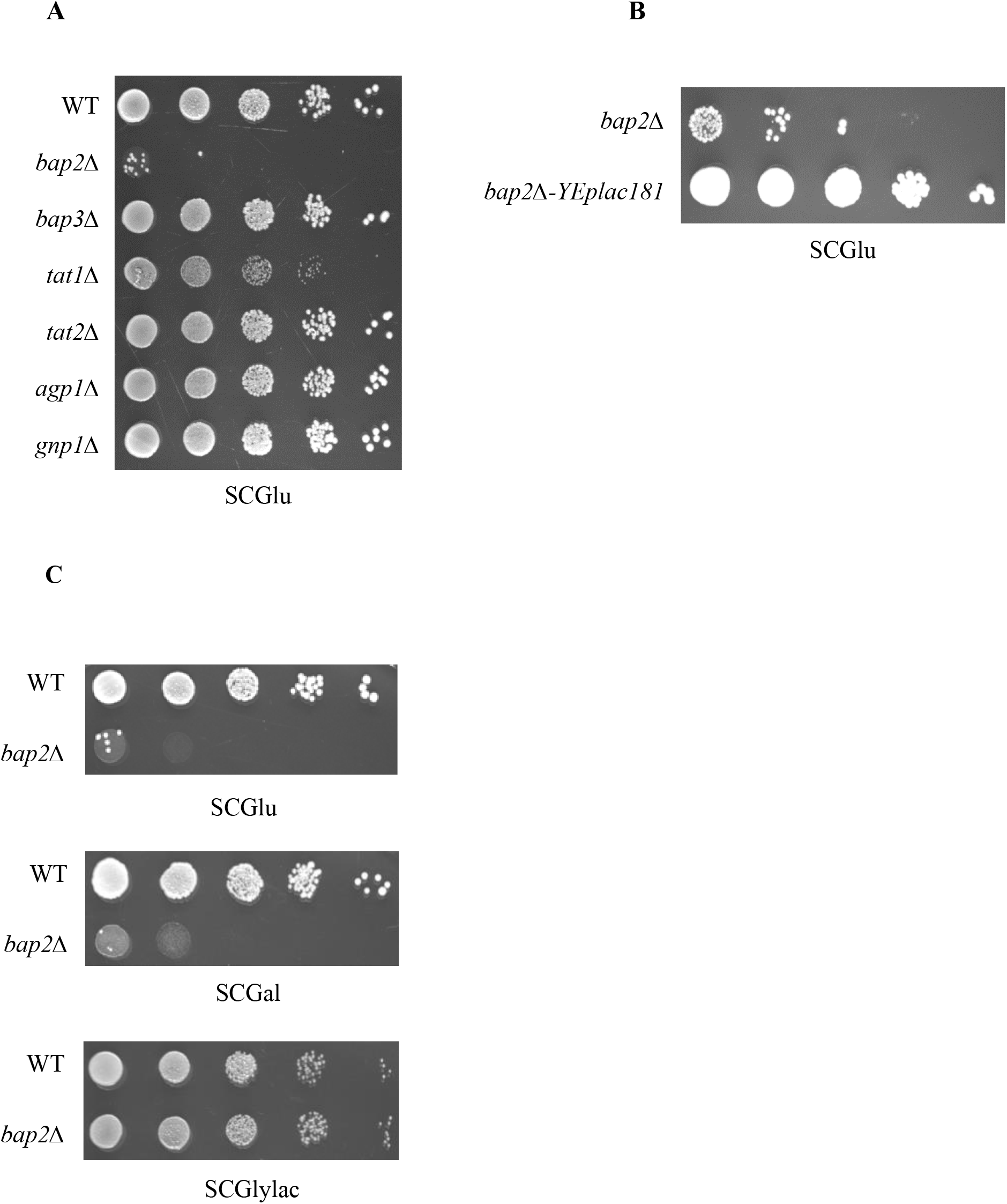
*BAP2* and *LEU2* are synthetically lethal in a carbon source dependent manner. (A) WT, *bap2*Δ, *bap3*Δ, *tat1*Δ, *tat2*Δ, *agp1*Δ and *gnp1*Δ strains grown in SCGlylac broth were spotted on SCGlu medium. (B) *bap2Δ* strain and its transformant carrying YEplac181, a *LEU2* plasmid grown in SCGlylac or SLeu^-^Glylac broth were spotted on SCGlu medium. (C) WT and *bap2*Δ strains grown in SCGlylac broth were spotted on SCGlu, SCGal and SCGlylac media. Images were captured and analyzed after incubating the petri plates for 2-3 days at 30°C.

The failure of the strains to import leucine in modified SCGlu medium was intriguing, in view of the presence of several leucine permeases such as Bap2, Bap3, Tat1, Tat2, Agp1, Gnp1 etc. We sought to ascertain the importance of these permeases in growth. We studied the growth of strains lacking leucine permeases and found that loss of *BAP2* could establish the leucine import deficient growth defect in the routine SCGlu medium (Fig.1A). The growth of *bap2Δ* strain was restored by *LEU2* complementation (Fig.1B). Among other permeases loss of *TAT1* conferred a mild growth defect. Interestingly, the growth defect vanished when the carbon source was swapped to glycerol and lactate (SCGlylac-<u>S</u>ynthetic <u>C</u>omplete <u>Gly</u>cerol and <u>lac</u>tate medium) but not galactose (SCGal-<u>S</u>ynthetic <u>C</u>omplete <u>Gal</u>actose medium) (Fig.1C).

We report for the first time that *BAP2* and *LEU2* are synthetically lethal in SCGlu/Gal medium. But why would loss of *BAP2*, confer growth retardation when other leucine permeases are also expressed in SCGlu medium (Forsberg et al., 2001)? Perhaps, *BAP2* is the predominant leucine transporter and its loss amounts to growth retardation.

### Growth defect of bap2Δ strain is overcome by loss of BUL1

We wondered if the growth defect be overcome by ways other than complementation by *LEU2*. In a tryptophan import deficient mutant, loss of *BUL1*, an arrestin restores import by stabilizing *TAT2*, a permease which imports Tryptophan, Tyrosine, Phenylalanine and Leucine etc. in plasma membrane (Umebayashi and Nakano, 2003). Actually, loss of *BUL1*, in the *bap2Δ* strain did restore growth normalcy in SCGlu and SCGal media (Fig.2A). But the growth of *bap2Δbul1Δ* strain was not dependent on *TAT2, GNP1* or *AGP1*, but *BAP3* (Fig.S1A,B). We speculated that loss of *BUL1*, led to *BAP3* accumulation in plasma membrane.

**Figure 2.**
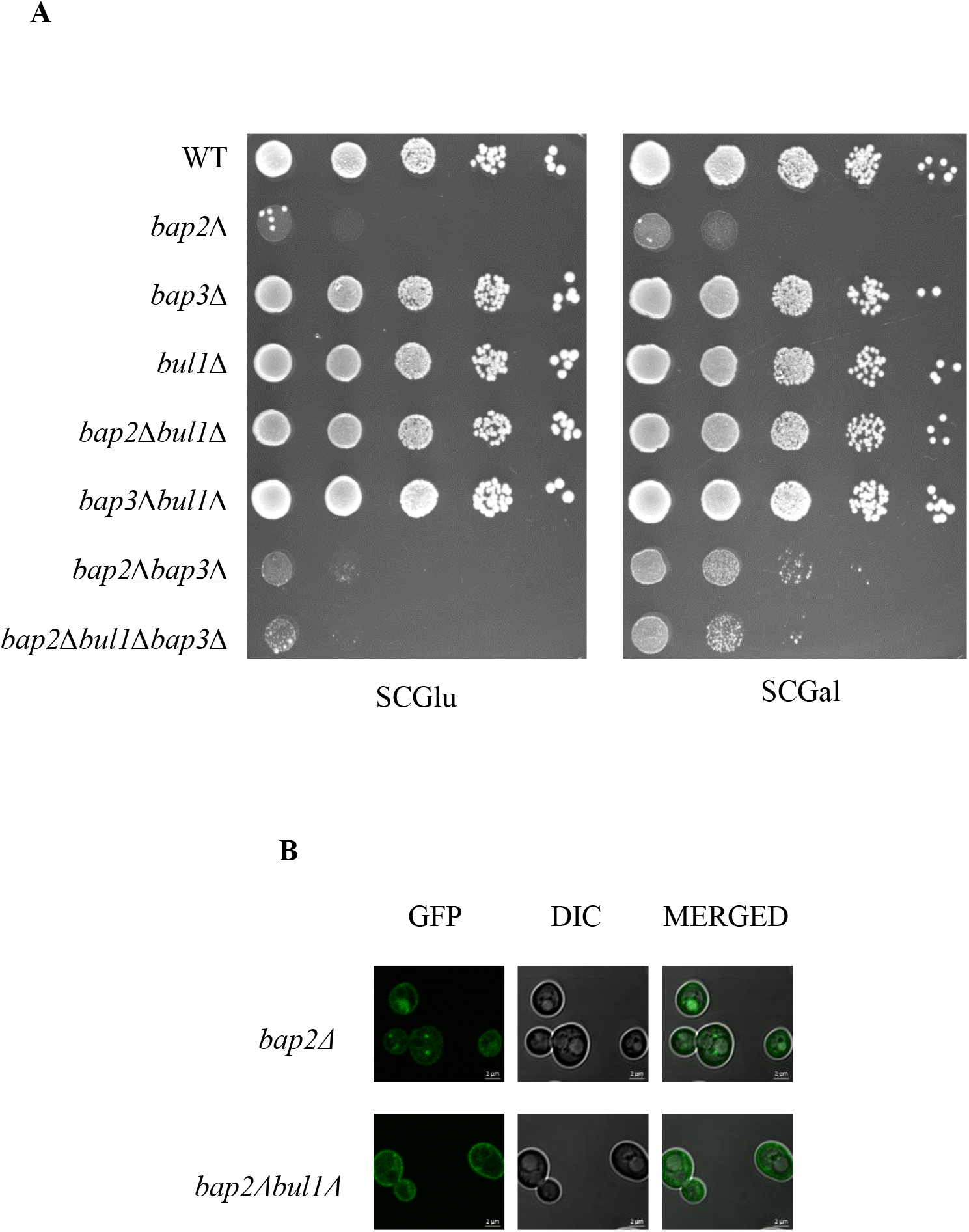
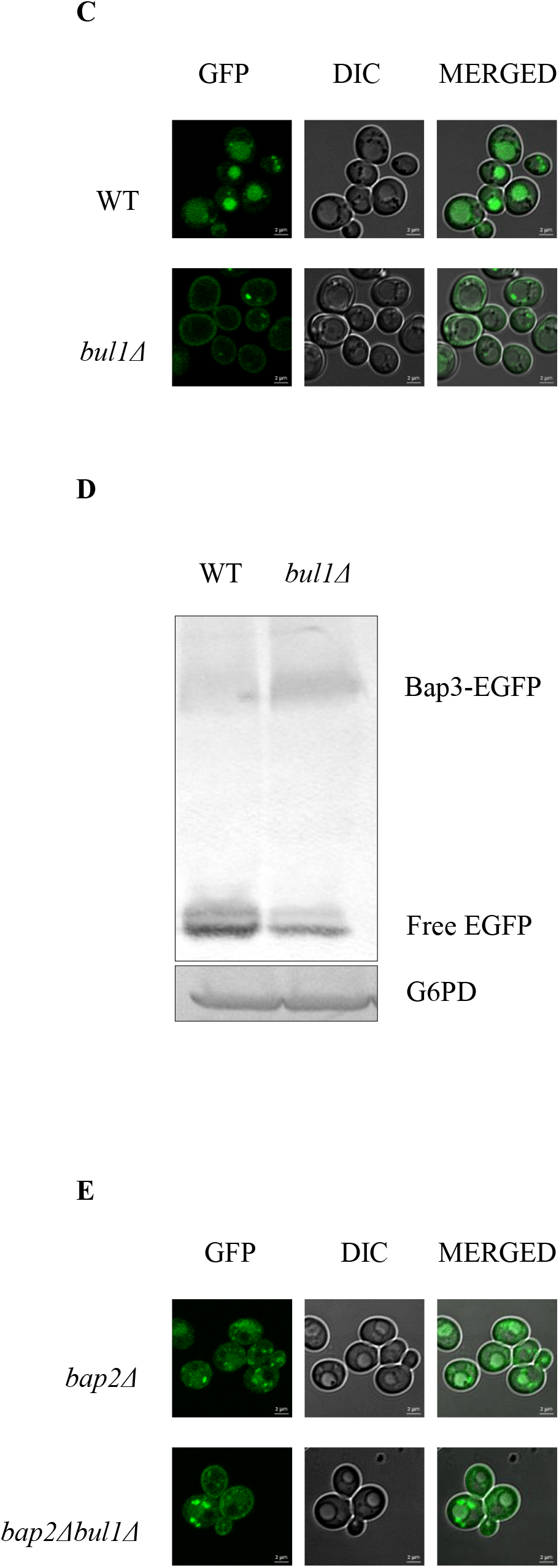
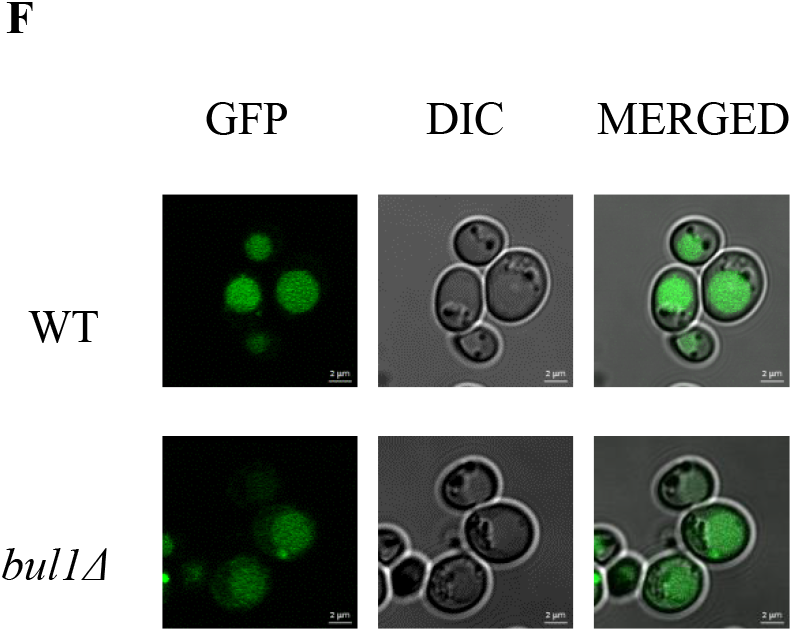
Loss of *BUL1* suppresses *bap2*Δ growth defect, by stabilizing Bap3 in the plasma membrane. (A) WT, *bap2*Δ, *bap3*Δ, *bul1*Δ, *bap2*Δ*bul1*Δ, *bap3*Δ*bul1*Δ, *bap2*Δ*bap3*Δ and *bap2*Δ*bul1*Δ*bap3*Δ strains grown in SCGlylac broth were spotted on SCGlu and SCGal media. Images were captured and analyzed after incubating the petri plates for 2-3 days at 30°C. (B) Localization of endogenously EGFP tagged *BAP3* in *bap2*Δ and *bap2*Δ*bul1*Δ strains grown in SCGlylac broth and subcultured in SCGlu broth upto mid logarithmic phase. Scale-2μM. (C) Localization of endogenously EGFP tagged *BAP3* in WT and *bul1*Δ strains grown in SCGlylac broth and subcultured in SCGlu broth upto mid logarithmic phase. Scale-2μM. (D) Immunoblotting of total cell lysates of endogenously EGFP tagged *BAP3* in WT and *bul1*Δ strains, grown as mentioned in C. Anti-GFP and anti-G6PD antibodies (for loading control) were used for immunoblotting. Free EGFP emanates from vacuolar degradation of Bap3-EGFP chimera. (E) Localization of endogenously EGFP tagged *BAP3* in *bap2*Δ and *bap2*Δ*bul1*Δ strains grown in SCGlylac broth and subcultured in SCGal broth upto mid logarithmic phase. Scale-2μM. (F) Localization of endogenously EGFP tagged *BAP3* in WT and *bul1*Δ strains grown in SCGlylac broth and subcultured in SCGal broth upto mid logarithmic phase. Scale-2μM.

As expected Bap3 accumulation was enriched in the periphery of the *bap2Δbul1Δ* strain than the *bap2Δ* strain (Fig.2B). Contrastingly, vacuolar Bap3 distribution was enriched in a *bap2Δ* strain than the *bap2Δbul1Δ* strain. This pattern of Bap3 distribution was more prominent while comparing *bul1Δ* and WT strains (Fig.2C). This observation shows that irrespective of the presence or absence *BAP2*, Bul1 downregulates Bap3. When blotted, membranous Bap3-EGFP population was enriched in *bul1Δ* strain than the WT (Fig.2D).

These results show that loss of *BUL1*, results in *BAP3* accumulation in cell membrane. Going by its canonical role, we surmise that *BUL1* is responsible for downregulation of *BAP3* in plasma membrane through Rsp5 mediated endocytosis. Unlike SCGlu medium, we did not observe any discernible difference in Bap3 distribution between *bap2Δ* and *bap2Δbul1Δ* or WT and *bul1Δ* strains grown in SCGal medium (Fig.2E,F).

### Loss of BUL1 empowers WT to grow in an extremely low leucine environment

We wondered how the *bap2Δ* strain would grow in a medium devoid of amino acids other than auxotrophic needs. Hence we chose SD medium (Synthetic Minimal Dextrose) for this study (Fig.3A). The *bap2Δ* strain grew well as WT in SD75 medium (75μM leucine), but ceased to grow at SD25 medium (25μM leucine). The growth was restored through loss of *BUL1*, and required *BAP3*. At SD5 medium (5μM leucine), even WT ceased to grow. Once again, loss of *BUL1* restored WT its growth which required *BAP2* not *BAP3*.

**Figure 3.**
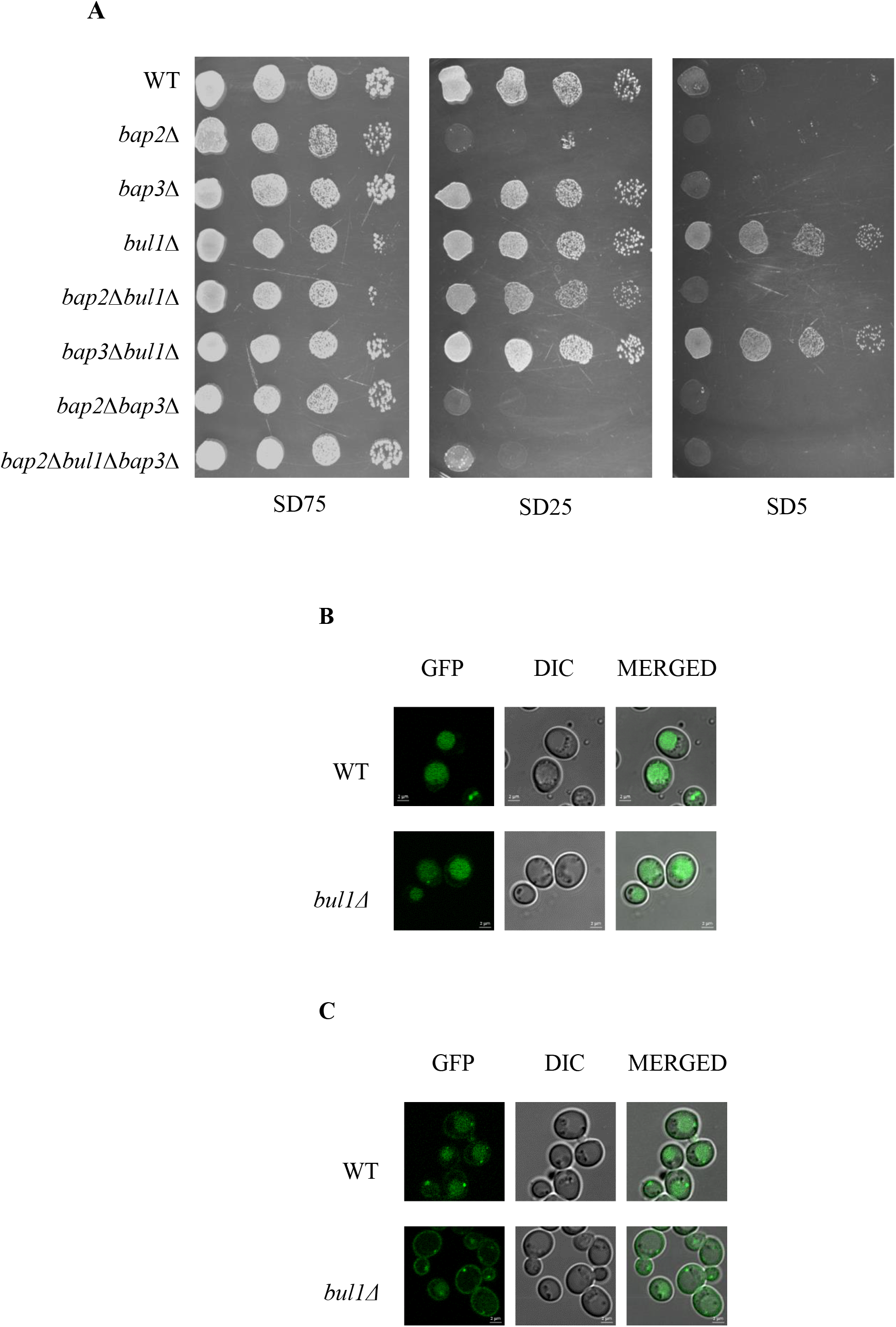
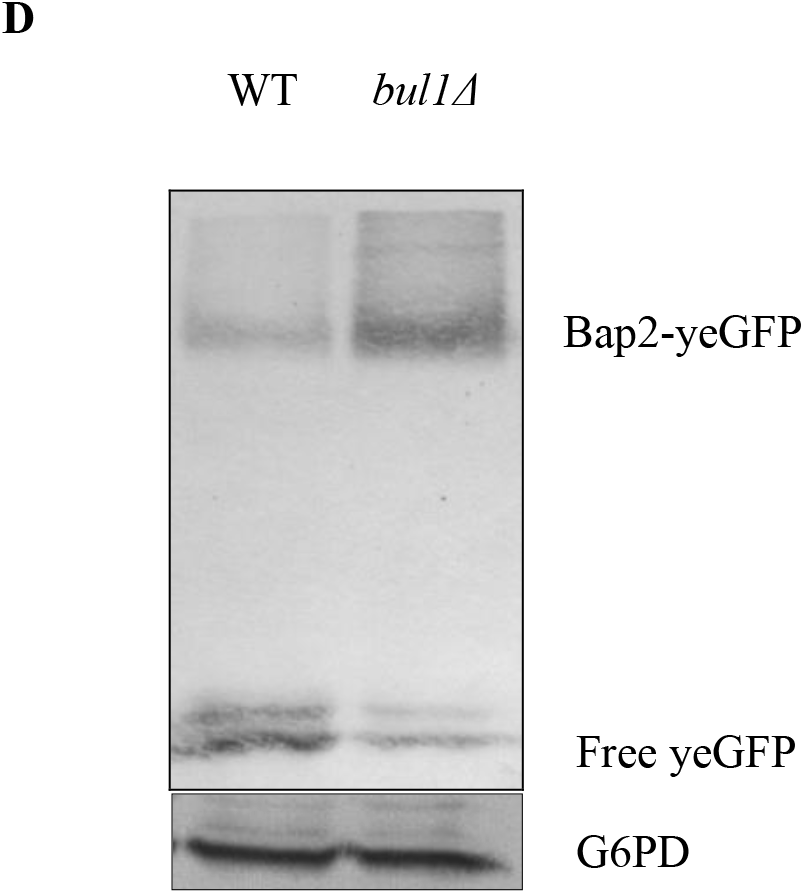
Loss of *BUL1* enables WT to grow in a medium with scarce leucine, through stabilization of Bap2 in the plasma membrane. (A) WT, *bap2*Δ, *bap3*Δ, *bul1*Δ, *bap2*Δ*bul1*Δ, *bap3*Δ*bul1*Δ, *bap2*Δ*bap3*Δ and *bap2*Δ*bul1*Δ*bap3*Δ strains grown in SCGlylac broth were spotted on SD75, SD25 and SD5 media. Images were captured and analyzed after incubating the petri plates for 2-3 days at 30°C. (B) Localization of endogenously EGFP tagged *BAP3* in WT and *bul1*Δ strains grown in SCGlylac broth and subcultured in SD25 broth upto mid logarithmic phase. Scale-2μM. (C) Localization of endogenously yeGFP tagged *BAP2* in WT and *bul1Δ* strains grown in SCGlylac broth and subcultured in SD25 broth upto mid logarithmic phase. Scale-2μM. (D) Immunoblotting of total cell lysates of endogenously yeGFP tagged *BAP2* in WT and *bul1*Δ strains, grown as mentioned in C. Anti-GFP and anti-G6PD antibodies (for loading control) were used for immunoblotting. Free EGFP emanates from vacuolar degradation of Bap2-yeGFP chimera.

These results show that *BAP2* and *LEU2* are synthetic lethal when (1) surplus of competing amino acids are present (SCGlu/Gal) or (2) leucine is scarce (SD). For the *bap2Δ* strain at SD75, absence of competing amino acids favors seamless leucine import through other permeases. This holds true for a *bap2Δbap3Δ* strain also. But in SD25, other permeases do not sustain the growth of *bap2Δ* strain, unless Bap3 accumulates at plasma membrane through loss of *BUL1*. In SD5, where leucine is very scarce, even WT doesn’t grow unless *BAP2* accumulates at plasma membrane due to loss of *BUL1*. We surmise that Bul1 downregulates both Bap2 and Bap3 at plasma membrane.

The *bul1Δ* strain displayed slightly increased in plasma membrane Bap3 localization, than the WT (Fig.3B). Though visible difference in vacuolar distribution was not observed. Unlike Bap3, Bap2 was enriched in plasma membrane of the *bul1Δ* strain than the WT and the vacuolar distribution was quite opposite (Fig.3C). For Fig.3C, we used SD25 broth as growing of cells in SD broth was extremely difficult. Immunoblotting results showed a larger Bap2-yeGFP and lesser free GFP population in the *bul1Δ* strain than the WT (Fig.3D). Our attempts to perform immunoblotting of Bap3-EGFP was not quite successful probably due to much reduced expression of Bap3-EGFP. These results indicate that loss of *BUL1*, leads to accumulation of Bap2 and Bap3 in plasma membrane possibly due to reduced downregulation.

### BAP2 is the major leucine transporter in SCGlu medium

As we suppose *BAP2* as the major leucine transporter in SCGlu medium, we tried to know how much leucine does *BAP2* import, how loss of *BUL1* affects the import in *bap2Δ*strain? etc. We measured the import of C14-L-Leucine in SCGlu medium for strains as in (Fig.4A). As strains like *bap2Δ* doesn’t grow in SCGlu medium, all the strains were made prototrophic for leucine to facilitate their growth. We assumed that the leucine prototrophy doesn’t affect the import by the permeases and their regulation by *BUL1*.

**Figure 4.**
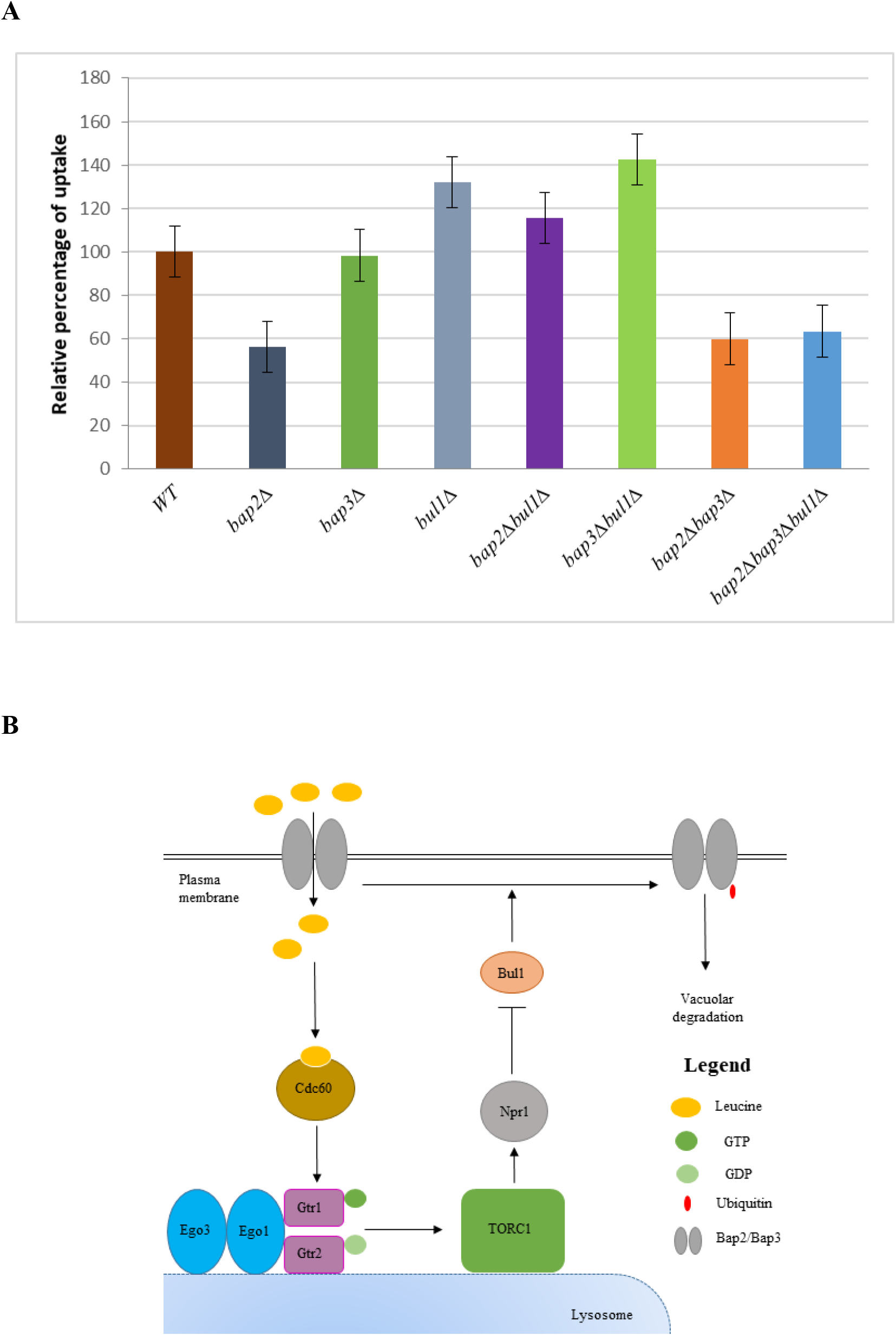
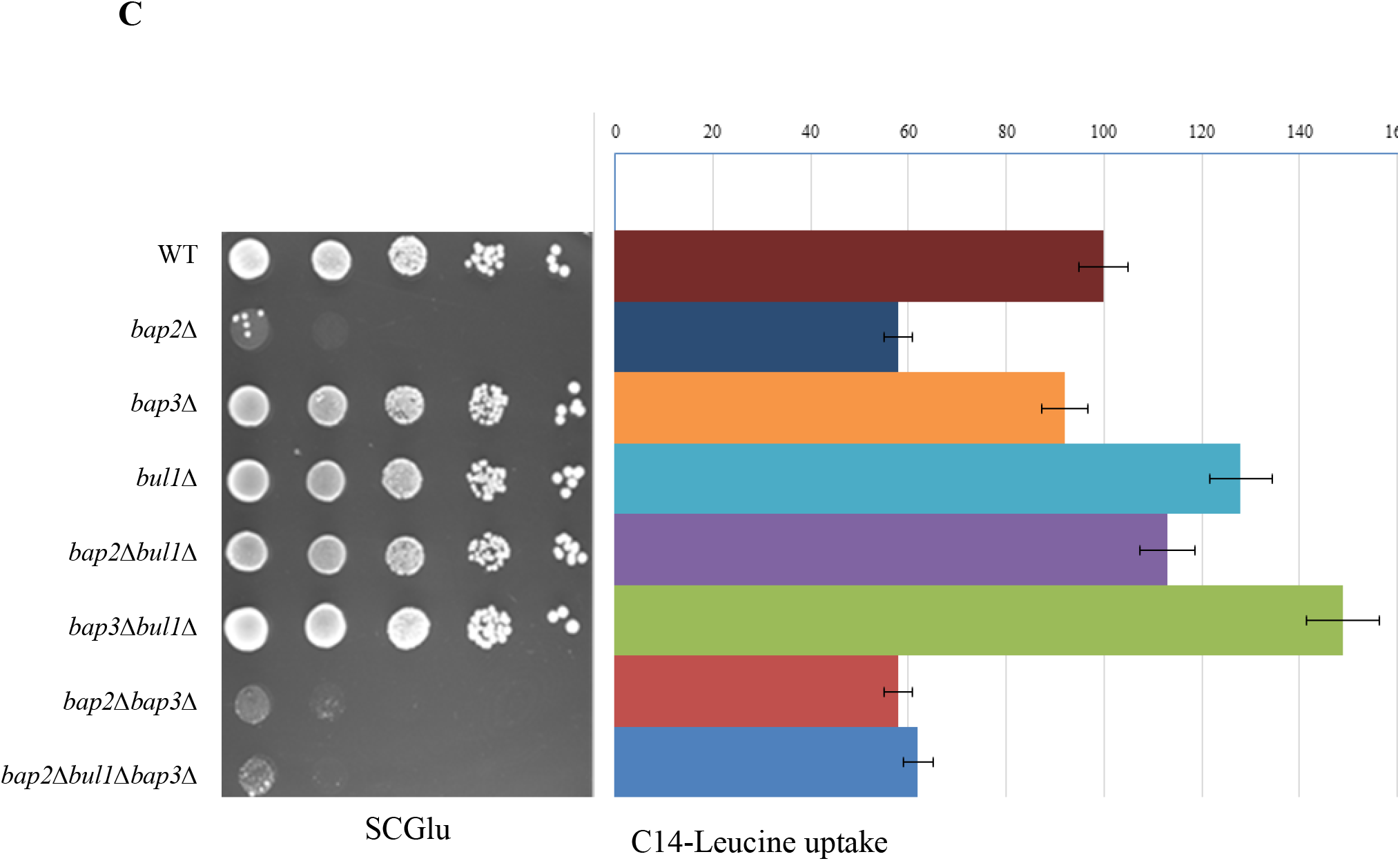
Leucine availability is crucial for growth. (A) WT, *bap2*Δ, *bap3*Δ, *bul1*Δ, *bap2*Δ*bul1*Δ, *bap3*Δ*bul1*Δ, *bap2*Δ*bap3*Δ and *bap2*Δ*bul1*Δ*bap3*Δ transformants carrying YEplac181, a *LEU2* plasmid were assayed for the uptake of C14 labelled L-Leucine. Uptake was calculated as picomoles /0.8×10^6^cells/min and expressed as mean relative percentage ± SE, considering uptake of WT as 100%. (B) Schematic representation of the proposed TORC1-Leucine feedback loop representing how leucine regulates TORC1 activity. (C) Correlation of growth phenotype of strains grown on SCGlu medium as shown in Fig.2A and their relative leucine uptake as shown in Fig.4A.

Assuming WT import capacity as 100%, *bap3Δ, bap2Δ* and *bap2Δbap3Δ* strains displayed 10%, 40% and 40% drop respectively. This implied *BAP2* as the major and *BAP3* as a minor leucine transporters in SCGlu medium. In *bap2Δ* strain, loss of *BUL1* doubled its import, which vanished on loss of *BAP3*. Interestingly, loss of *BUL1* in the *bap3Δ* strain augmented its import by 50%. This was due to apparent accumulation of *BAP2* at the plasma membrane, as loss of *BAP2* not only eliminated this effect but also reduced its import alike to that of a *bap2Δ* strain. Loss of *BUL1*, in WT increased the import by 30%, possibly due to plasma membrane accumulation of both *BAP2* and *BAP3*. Import data for *bap2Δ* and *bap3Δ* strains agree with earlier reports (Grauslund et al., 1995; Didion et al., 1998).

### Loss of BUL2 does not suppress the growth defect of bap2Δ strain

*BUL2* and *BUL1* are paralogues, together they downregulate many permeases (Yashiroda et al., 1998; O’Donnell and Schmidt, 2019). WT strain used in this study is naturally *bul2Δ* as it harbors a loss of function allele of *BUL2* (*BUL2^F883L^*) (Kwan et al., 2011). Perhaps the suppression of the *bap2Δ* growth defect through accumulation of Bap3 at plasma membrane is due to the collective loss of *BUL1* and *BUL2*. In such case, we suppose that the expression of a functional *BUL2* in a *bap2Δbul1Δ* strain (precisely *bap2Δbul1Δbul2Δ*) shall favor downregulation of Bap3 and confer back the growth defect at least partially if not fully.

To check this idea, the *bap2Δbul1Δ* strain was complemented with a functional *BUL2* allele from W303-1A strain background alongside *BUL1* as a positive control (Fig.S2). But it was not *BUL2* but *BUL1* to confer the growth defect back. We believe that *BUL2* does not downregulate Bap3 at plasma membrane and the suppression is solely due to loss of *BUL1* but not *BUL2*. The functionality of the *BUL2* allele was verified as it successfully restored the growth of the *bul1Δ* strain (precisely *bul1*Δ*bul2*Δ) which suffered a severe growth defect at 37°C in SMTGlu (<u>S</u>ynthetic <u>M</u>inimal <u>T</u>yrosine <u>Glu</u>cose medium) (Fig.S3).

## Discussion

Organisms auxotrophic for nutrients rely on import from extracellular medium. Though *Saccharomyces cerevisiae* is naturally prototrophic for all amino acids, lab strains are usually auxotrophic for afew amino acids. Strains auxotrophic for leucine, tyrosine and tryptophan have difficulty in growth due to their poor import of corresponding amino acids from the media with surplus amino acids (Cohen and Engelberg, 2007).

In this study, in an otherwise normally growing WT strain (*leu2Δ*), loss of *BAP2* confers a growth defect due to poor import of leucine. The growth defect was suppressed by accumulation of Bap3 at plasma membrane due to loss of an arrestin, *BUL1*. Also in a *BAP2*^+^ strain, loss of *BUL1* leads to plasma membrane accumulation of Bap2. With these results we propose Bul1 as a post translational regulator to regulate the abundance of Bap3 and Bap2 at plasma membrane possibly through ubiquitin conjugated endocytosis. This finding joins the long list of transporters regulated by Bul1.

SPS (Ssy1-Ptr3-Ssy5) is an amino acid sensor present in plasma membrane. *SSY1* or *PTR3* and *LEU2* are synthetically lethal in SCGlu medium, overcome by loss of *BUL1* (Forsberg et al., 2001). This study finds that the pair *BAP2* and *LEU2* alone is sufficient to confer synthetic lethality in SCGlu and other media too. In presence of extracellular amino acids, SPS amino acid sensor prompts transcription of permeases *AGP1, GNP1, BAP2, BAP3, TAT1* and *TAT2*, all of which can import leucine at varying rates (Forsberg et al., 2001). We suppose *BAP2* as the major importer of leucine as loss of *BAP2* alone causes the growth defect and 40% drop in import. Galactose and glycerol upsurge leucine import by tenfold than glucose (Peter et al., 2006), yet the growth defect persists in SCGal not in SCGlylac medium. Again it seems that *BAP2* is the major leucine importer in SCGal, while in SCGlylac other leucine permeases account for the hike. These observations oblige to revisit the amino acid metabolism/uptake in terms of the Carbon source, instead the Nitrogen source or amino acids as it has been customarily.

We speculate a signaling loop whereby leucine regulates its import (Fig.4B). Leucine binds Cdc60, a leucyl tRNA synthetase cum intracellular leucine sensor to activate TORC1, via EGO complex bound Rag GTPases Gtr1 and Gtr2 (Bonfils et al., 2012). Active TORC1 releases Bul1 from Npr1 kinase inhibition (Merhi and Andre, 2012). Bul1 promotes Bap2 and Bap3 endocytosis, causing drop in leucine import and intracellular levels, eventually reducing TORC1 activity.

Loss of *BUL2* reduces chronological lifespan (CLS) and prolongs replicative lifespan (RLS), due to suspected increased TORC1 activity, as a result of increased intracellular amino acid pool (Kwan et al., 2011). Then how the loss of *BUL1* would affect the lifespans? With our observations and fact that Leucine activates TORC1, we expect *BUL1* to have a greater bearing than *BUL2*. Contradicting the above, CLS of a *leu2Δ* strain is extended by leucine prototrophy or increasing leucine concentration in the medium (Alvers et al., 2009).

Being the most sought amino acid for translation, a slight shortage of leucine may harshly affect the growth. Adhering to this notion *bap2Δ, bap2Δbap3Δ* and *bap2Δbap3Δbul1Δ* strains that show 40% reduction in leucine import are the ones to suffer growth defect (Fig.4C). We think of two reasons to explain this scenario. The intracellular leucine concentration is below the threshold (1) to bind Cdc60 and activate TORC1. As a result growth ceases when TORC1 is not active. (2) to charge Cdc60 for translation.

## Supplementary figures

**Figure S1.**
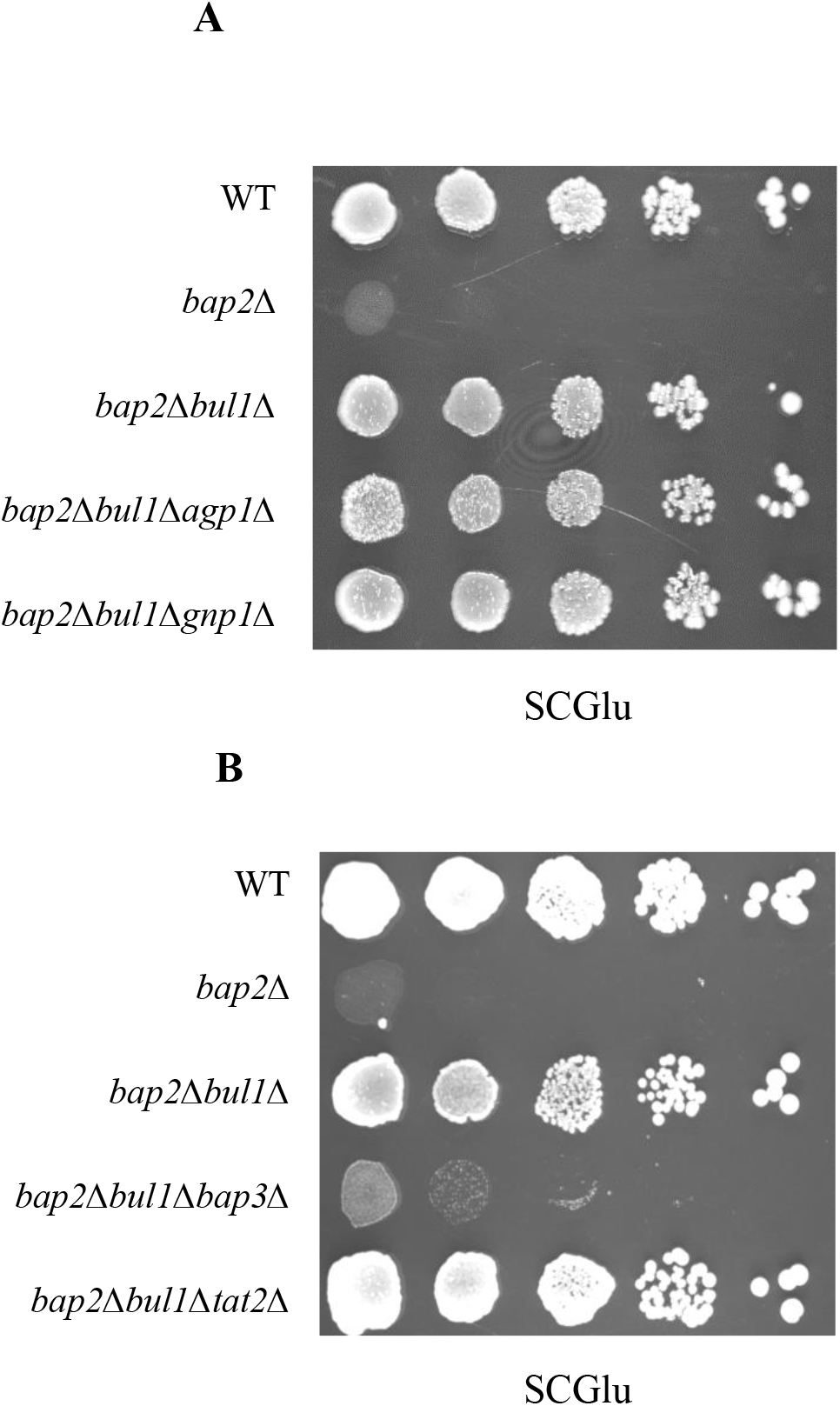
Loss of *BUL1* suppresses the *bap2*Δ growth defect, in a *BAP3* dependent manner. (A) WT, *bap2*Δ, *bap2*Δ*bul1*Δ, *bap2*Δ*bul1*Δ*agp1*Δ and *bap2*Δ*bul1*Δ*gnp1*Δ strains grown in SCGlylac broth were spotted on SCGlu medium. (B) WT, *bap2*Δ, *bap2*Δ*bul1*Δ, *bap2*Δ*bul1*Δ*bap3*Δ and *bap2*Δ*bul1*Δ*tat2*Δ strains grown in SCGlylac broth were spotted on SCGlu medium. Images were captured and analyzed after incubating the petri plates for 2-3 days at 30°C.

**Figure S2.**
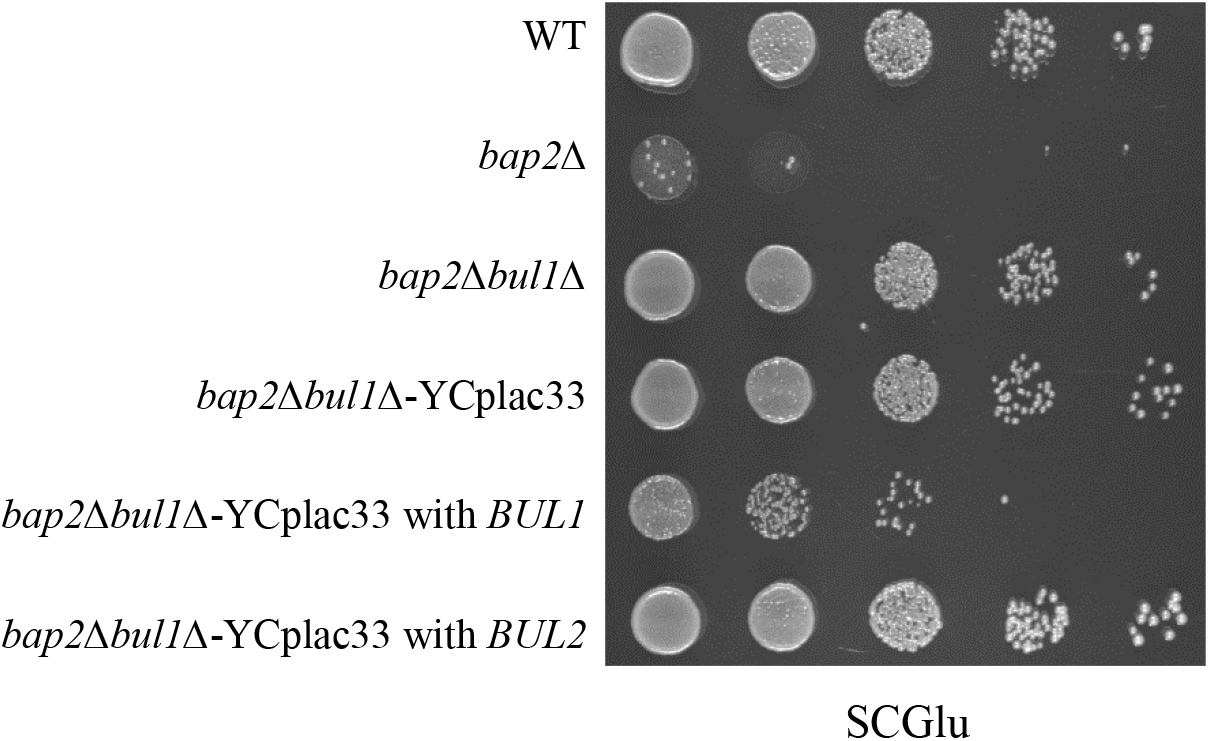
Complementation by *BUL1* not *BUL2* confers the growth defect back in the *bap2*Δ*bul1*Δ strain. WT, *bap2*Δ, *bap2*Δ*bul1*Δ strains and *bap2*Δ*bul1*Δ transformants carrying YCplac33 or YCplac33 with *BUL1* or *BUL2* grown in SCGlylac or SUra^-^Glylac broth were spotted for growth on SCGlu medium. Images were captured and analyzed after incubating the petri plates for 2-3 days at 30°C.

**Figure S3.**
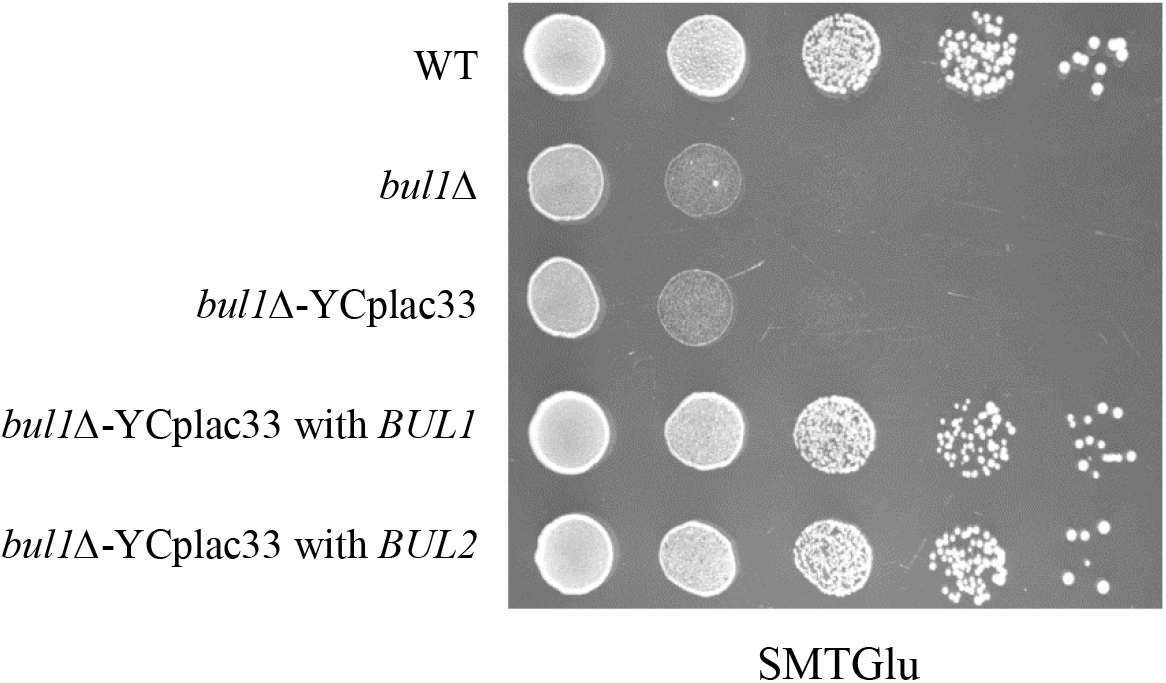
*BUL2* of W303-1A strain is functional. WT, *bul1*Δ strains and *bul1*Δ transformants carrying YCplac33 or YCplac33 with *BUL1/BUL2* grown in SCGlylac or SUra^-^Glylac broth were spotted for growth on SMTGlu medium. Images were captured and analyzed after incubating the petri plates for 2-3 days at 37°C.

## Materials and methods

### Yeast strains and plasmids

All yeast strains are of BY4741 background (ST.1). They were either purchased from Euroscarf or constructed. Yeast transformation was performed by LiAc-PEG method. Yeast genomic DNA was isolated by glass bead method. Genetic manipulations like disruption and tagging were performed by PCR based homologous recombination method. These manipulations were again confirmed by PCR. Primers used for the above procedures are listed (ST.3).

E.coli strain XL1 or DH5α were used for plasmid cloning and propagation. Plasmid isolation was done by alkaline lysis method and *E.Coli* transformation was done by PEG method. Plasmids used in the study are listed (ST.2). Plasmid carrying *BUL1* was constructed by cloning of 4020 bp SalI-EcoRI fragment of *BUL1* gene (1000 bp upstream of start codon, 2931 bp CDS and 89 bp downsream of stop codon), into SalI-EcoRI site of YCplac33. *BUL1* was amplified from the genomic DNA of WT BY4741. Plasmid carrying *BUL2*, was constructed by cloning of a 4019 bp SalI-SacI fragment of *BUL2* (1000 bp upstream of start codon, 2763 bp CDS, 250 bp downsream of stop codon, 6 bp of primer region), amplified from genomic DNA of WT W303-1A strain, into the SalI-SacI site of YCplac33.

#### Make

Primers (Sigma, BLOT solutions), Restriction enzymes, polymerases and ligases (Thermo scientific, NEB), PCR/Gel purification kits (Qiagen).

### Media recipe

#### Synthetic Complete Glucose or Synthetic Complete Galactose (SCGlu/SCGal)

0.17% Yeast nitrogen base, 0.5% ammonium sulfate, 0.05% amino acid mixture (ST4) and 2% glucose or galactose. **Synthetic Complete Glycerol plus lactate (SCGlylac)** – 0.17% Yeast nitrogen base, 0.5% ammonium sulfate, 0.05% amino acid mixture (ST4) and 3% (V/V) glycerol and 2% (V/V) of 40% lactate pH 5.7. **Synthetic Minimal Dextrose (SD)** – 0.17% Yeast nitrogen base, 0.5% ammonium sulfate, 2% glucose and 0.01% of auxotrophic supplements Uracil (89μM), Histidine (64μM), Methionine (67μM) and Leucine (76μM). SD75, SD25 and SD5 refer to SD with 75μM or 25μM or 5μM leucine. For solid media 2%agar was used. SUra^-^Glylac and SLeu^-^Glylac refers to SCGlylac lacking Uracil or Leucine respectively. **Synthetic Minimal Tyrosine Glucose (SMTGlu)** – 0.17% Yeast nitrogen base, 2% glucose, 0.01% of auxotrophic supplements and 2mM Tyrosine.

### Spotting assay (Growth assay)

Cells were grown in SCGlylac broth (SCGlylac broth lacking uracil or leucine for transformants) to mid log phase, harvested and washed with sterile distilled water and cell density was normalized to OD600 1.0. The normalized cell suspension was serially diluted to tenfold dilutions upto 10^-4^ dilution. 5μl from each dilution was spotted onto appropriate media and incubated at 30°C or 37°C for 2-3 days. Images of the petriplates were captured by UVITEC, CAMBRIDGE gel documentation system of GeNei.

### Protein extraction and immunoblotting

Protein was extracted by alkaline lysis method. Cells grown in suitable medium to mid log phase were harvested at 4°C and lysed with 0.5ml of lysis solution (0.2M NaOH, 0.2% mercaptoethanol) for 10 min on ice. Trichloroacetic acid was added to a final concentration of 5%, and incubated for another 10 min on ice. Samples were centrifuged for 5 min at 13000 x g for 5 min. Pellets were resuspended in 35 μl of dissociation buffer (4% SDS, 0.1M Tris-HCl pH 6.8, 4mM EDTA, 20% glycerol, 2% 2-mercaptoethanol, 0.02% Bromophenol blue) and mixed with 15 μl of 1M Tris base, and heated at 37°C for 10 min.

Total cell free extracts were resolved by SDS-PAGE and transferred to 0.45 micron nitrocellulose membrane. Mouse bi-clonal anti-GFP antibodies were used to probe GFP tagged proteins and chased by secondary anti-mouse goat antibody conjugated with alkaline phosphatase. Rabbit anti-G6PD antibody was used to probe G6PD, and chased down by secondary anti-rabbit mouse antibody conjugated with alkaline phosphatase. Blots were developed and images were captured by UVITEC, CAMBRIDGE gel documentation system of GeNei and analyzed by UVITEC1D software.

#### Make

Mouse bi-clonal anti-GFP antibody (Roche Diagnostics, Mix of 13.1 and 7.1 monoclonal antibodies), Rabbit anti-G6PD antibody (Santacruz Biotechnology) and Secondary anti-rabbit mouse antibody (Santacruz Biotechnology). Nitrocellulose membrane (Amersham Biosciences). SDS-PAGE and blotting transfer apparatus (GeNei).

### Fluorescence microscopy

Cells were grown in appropriate media to mid log phase were harvested, washed in sterile distilled water and resuspended in the same (To avoid auto fluorescence due to nutrient broth). Cells were readily analyzed for GFP fluorescence at room temperature under Zeiss Axio-Observer Z1 inverted motorized and computer-controlled laser scanning fluorescence microscope (Carl Zeiss, LSM 780) equipped with iPlan-Apochromat 100X oil immersion objective 1.4 NA and PMT detector camera. Argon+ laser was used to excite the fluorophore with excitation wavelength of 488nm and emission wavelength was 526nm. Images were captured and later processed by Zen 2012 blue acquisition software from Zeiss.

### Leucine uptake assay

Yeast cells were grown in SCGlylac broth lacking leucine; subcultured in SCGlu broth and grown upto mid log phase (OD_600_ 1-1.5). Cells were harvested, washed with SCGlu broth twice and resuspended in the same to 0.4×10^7^cells/ml. To 200μl of this suspension added was 2.8KBq (500 picomoles) of C14-labelled leucine. The suspension contained 188μM of cold leucine and 2.5μM of C14-Leucine. The cell suspensions were incubated in 30°C incubator cum shaker for 0 min, 60 min and 120 min time points. The C14-labelled leucine uptake was stopped by adding cold leucine of 100μl to a final leucine concentration of 12.6mM. The cells from the suspension were collected on a 0.45 micron nylon membrane (Amersham Biosciences) by suction. Cells were washed with ice cold water for 3 times and the membrane was dried for 10 minutes. The membrane with cells was counted for radioactive pulses for 1min/sample in Perkin Elmer Liquid scintillation analyzer. For each repetition biological replicates were taken and technical triplicates were taken for each strain for each time point. C14-Leucine with specific activity of 5.5GBq/mmole (150mci/mmole) was purchased from BRIT, India.

## Supplementary tables

**ST1.** Yeast strains.

ST1. Yeast strains.
| Strain | Genotype | Source |
| --- | --- | --- |
| WT | MAT a *his3Δ1 leu2Δ0 met15Δ0 ura3Δ0* | Euroscarf |
| *bap2Δ* | MAT a *his3Δ1 leu2Δ0 met15Δ0 ura3Δ0 bap2Δ::KanMX* | This study |
| *bap3Δ* | MAT a *his3Δ1 leu2Δ0 met15Δ0 ura3Δ0 bap3Δ::KanMX* | This study |
| *tat1Δ* | MAT a *his3Δ1 leu2Δ0 met15Δ0 ura3Δ0 tat1Δ::KanMX* | This study |
| *tat2Δ* | MAT a *his3Δ1 leu2Δ0 met15Δ0 ura3Δ0 tat2Δ::KanMX* | This study |
| *agp1Δ* | MAT a *his3Δ1 leu2Δ0 met15Δ0 ura3Δ0 agp1Δ::KanMX* | This study |
| *gnp1Δ* | MAT a *his3Δ1 leu2Δ0 met15Δ0 ura3Δ0 gnp1Δ::KanMX* | This study |
| *bul1Δ* | MAT a *his3Δ1 leu2Δ0 met15Δ0 ura3Δ0 bul1Δ::hphMX* | This study |
| *bap2Δbul1Δ* | MAT a *his3Δ1 leu2Δ0 met15Δ0 ura3Δ0 bap2Δ::KanMX bul1Δ::hphMX* | This study |
| *bap3Δbul1Δ* | MAT a *his3Δ1 leu2Δ0 met15Δ0 ura3Δ0 bap3Δ::Sphis5 bul1Δ::hphMX* | This study |
| *bap2Δbap3Δ* | MAT a *his3Δ1 leu2Δ0 met15Δ0 ura3Δ0 bap2Δ::KanMX bap3Δ::Sphis5* | This study |
| *bap2Δbul1Δbap3Δ* | MAT a *his3Δ1 leu2Δ0 met15Δ0 ura3Δ0 bap2Δ::KanMX bul1Δ::hphMX bap3Δ::Sphis5* | This study |
| *bap2Δbul1Δtat2* | MAT a *his3Δ1 leu2Δ0 met15Δ0 ura3Δ0 bap2Δ::KanMX bul1Δ::hphMX tat2Δ::KIURA3* | This study |
| *bap2Δbul1Δagp1Δ* | MAT a *his3Δ1 leu2Δ0 met15Δ0 ura3Δ0 bap2Δ::KanMX bul1Δ::hphMX agp1Δ::KIURA3* | This study |
| *bap2Δbul1Δgnp1Δ* | MAT a *his3Δ1 leu2Δ0 met15Δ0 ura3Δ0 bap2Δ::KanMX bul1Δ::hphMX gnp1Δ::KIURA3* | This study |
| *bap2ΔBAP3-EGFP* | MAT a *his3Δ1 leu2Δ0 met15Δ0 ura3Δ0 bap2Δ::KanMX BAP3::EGFP-Sphis5* | This study |
| *bap2Δbul1ΔBAP3-EGFP* | MAT a *his3Δ1 leu2Δ0 met15Δ0 ura3Δ0 bap2Δ::KanMX bul1Δ::hphMX BAP3::EGFP-Sphis5* | This study |
| *BAP3-EGFP* | MAT a *his3Δ1 leu2Δ0 met15Δ0 ura3Δ0 BAP3::EGFP-Sphis5* | This study |
| *bul1ΔBAP3-EGFP* | MAT a *his3Δ1 leu2Δ0 met15Δ0 ura3Δ0 bul1Δ::hphMX BAP3::EGFP-Sphis5* | This study |
| *BAP2-yeGFP* | MAT a *his3Δ1 leu2Δ0 met15Δ0 ura3Δ0 BAP2::yeGFP-hphMX* | This study |
| *bul1ΔBAP2-yeGFP* | MAT a *his3Δ1 leu2Δ0 met15Δ0 ura3Δ0 bul1Δ::kanMX BAP2::yeGFP-hphMX* | This study |

**ST2.** Plasmids.

ST2. Plasmids.
| Plasmid | Description | Source |
| --- | --- | --- |
| YCplac33 | Yeast centromeric shuttle vector with *AMP<sup>R</sup>* and *URA3* markers | Lab stock |
| YCplac33-*BUL1* | Yeast centromeric shuttle vector with *AMP<sup>R</sup>* and *URA3* markers carrying *BUL1* | This study |
| YCplac33-*BUL2* | Yeast centromeric shuttle vector with *AMP<sup>R</sup>* and *URA3* marker carrying *BUL2* | This study |
| YEplac181 | Yeast episomal shuttle vector with *AMP<sup>R</sup>* and *LEU2* markers | Lab stock |

**ST3.**
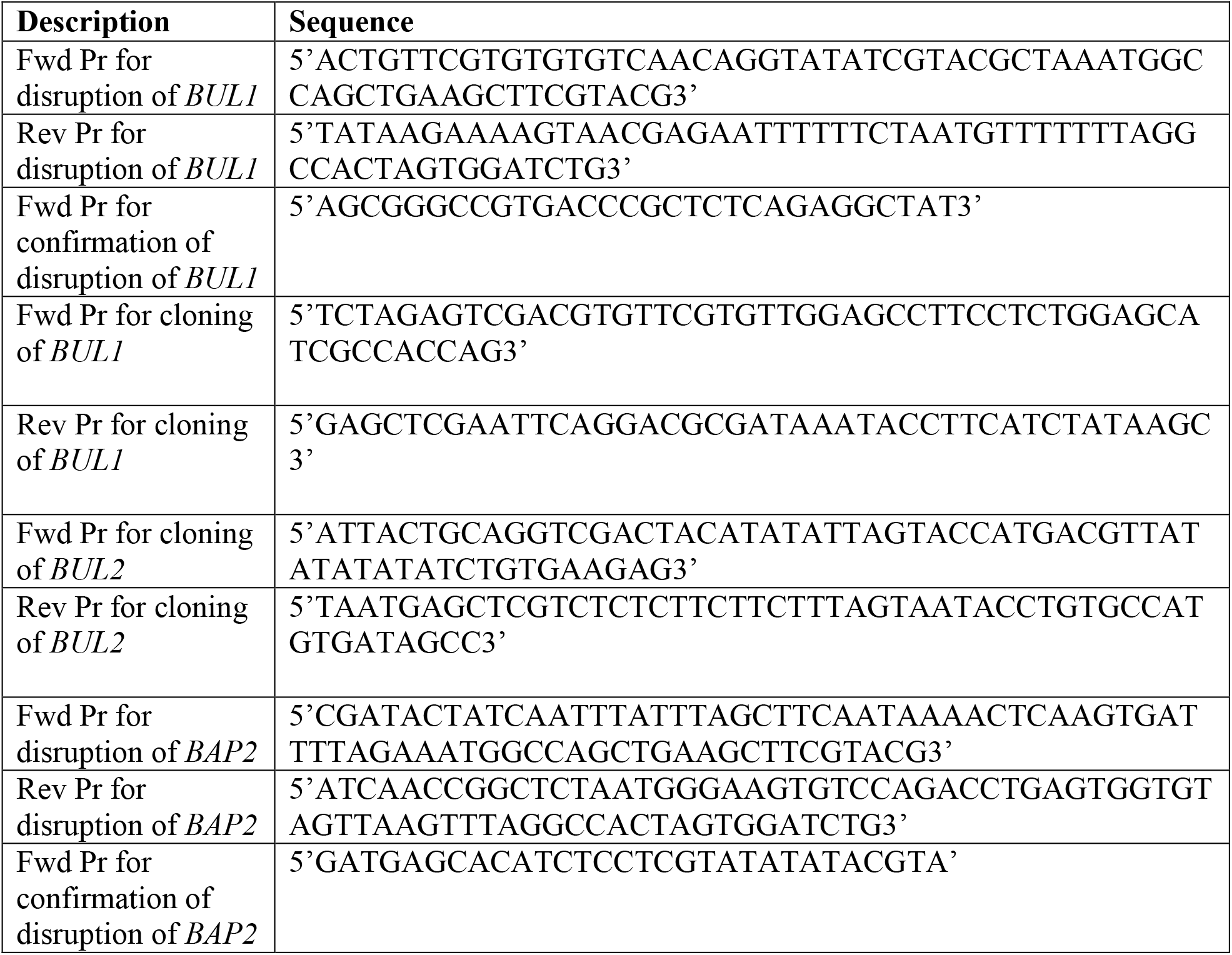
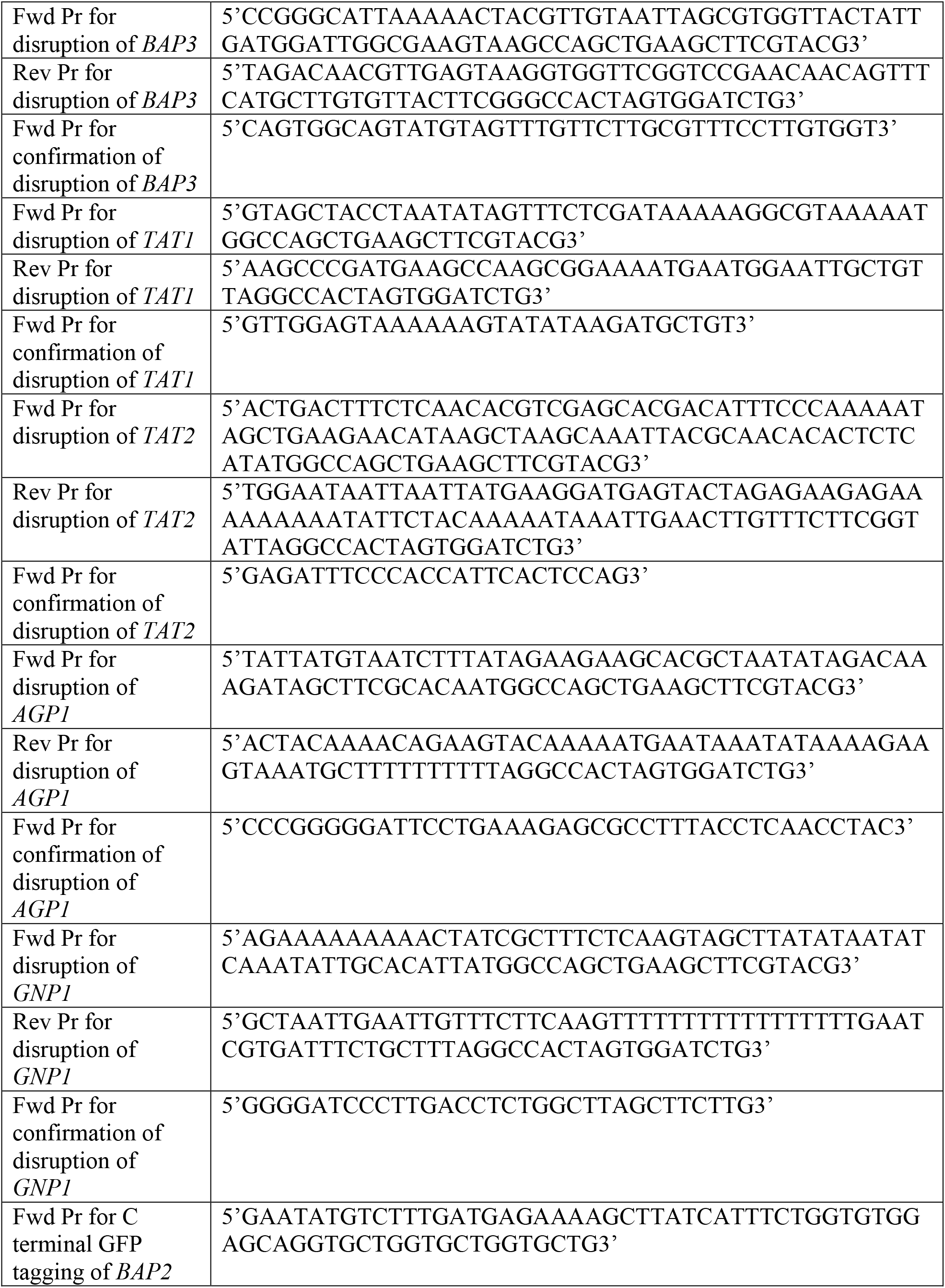
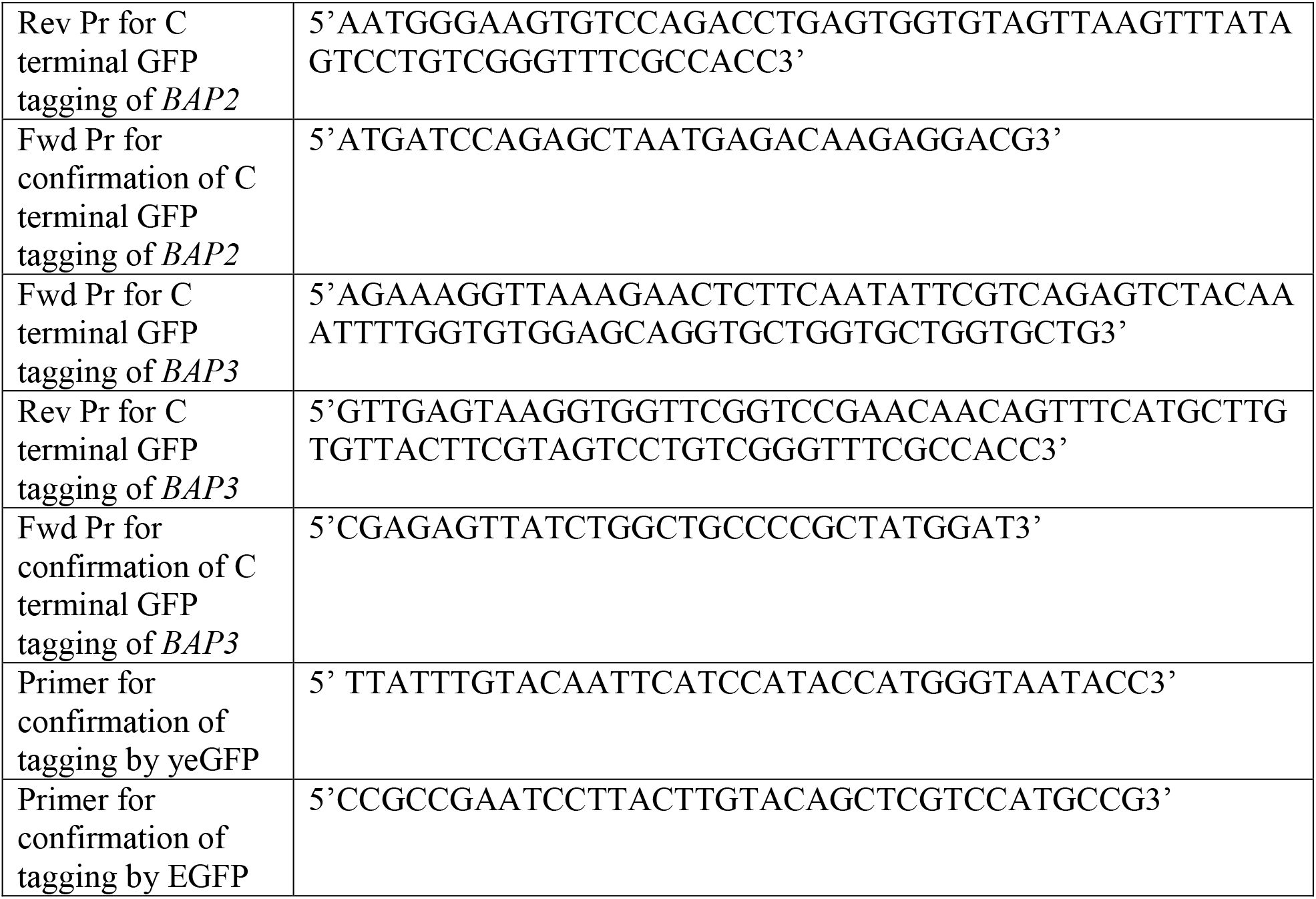
Primers.

## Acknowledgements

We are very grateful to Department of Science and Technology, INDIA, for funding the project (P13DST024).

## Author contribution

MS and PJB conceived ideas and designed the experiments. MS performed the experiments. MS and PJB analyzed the results. MS and PJB wrote the manuscript.

## Data availability statement

Data supporting the findings of this study are available, upon request.

